# Orchestrated Restructuring Events During Secretory Granule Maturation Mediate Intragranular Cargo Segregation

**DOI:** 10.1101/2021.08.16.456250

**Authors:** Zulfeqhar A. Syed, Liping Zhang, Duy T. Tran, Christopher K. E. Bleck, Kelly G. Ten Hagen

## Abstract

Regulated secretion is an essential process where proteins are packaged into membranous secretory vesicles. However, the details of cargo packaging and secretory granule maturation are largely unknown. Here, we demonstrate that multiple distinct proteins undergo intragranular restructuring during secretory granule maturation in vivo, resulting in spatial segregation of distinct protein components within the same granule. Furthermore, through a combination of genetics and multimodality imaging, we demonstrate the molecular identity of each distinct intragranular structure. We further identify temporally-regulated genes that are essential for the restructuring events, including those controlling pH (*Vha16-1*), Cl^−^ ions (*Clic* and *ClC-c*) and Ca^2+^ ions (*fwe*). Finally, we show that altered cargo glycosylation influences dimensions of these structures, thereby affecting secretory granule morphology. This study elucidates key steps and factors involved in intragranular, rather than intergranular segregation of cargo through regulated restructuring events during secretory granule maturation. Understanding how multiple distinct proteins are efficiently packaged into and secreted from the same secretory granule may provide insight into diseases resulting from defects in secretion.

## INTRODUCTION

Regulated secretion is an essential process where proteins are packaged into membranous secretory granules that then await a signal to deliver their contents to the extracellular space. While diverse cells and tissues undergo regulated secretion, the details of how specific proteins are transported through the secretory apparatus, appropriately modified and then subsequently targeted to and packaged within secretory granules remain largely unknown. Two basic models have been proposed where cargo is sorted into secretory granules either by “sorting for entry” or by “selective retention” (1), along with other models of unconventional protein secretion (2). In the bag cells of *Aplysia californica*, pro-egg laying hormone undergoes endoproteolytic cleavage in the Golgi complex to yield two peptides which are then sorted into two distinct vesicles and transported to different sites in the cell (3–5). Likewise, stratified squamous epithelia produce lamellar granules (LG) that contain distinct components that are independently trafficked to the cell surface (6). In contrast, in neuronal large dense-core vesicles (LDCVs), multiple components are heterogeneously distributed in a secretory granule and co-transported (7–13). When many proteins do reside within the same secretory granule, questions arise regarding how these diverse proteins are organized and segregated so that they may be appropriately secreted upon fusion of the granule with the plasma membrane of the cell. The issues of protein transport, compaction, segregation and secretion are of particular importance for large extracellular matrix proteins, many of which undergo regulated hydration and expansion as they are secreted from specialized cells (14–19).

Mucins are a family of large, highly O-glycosylated apical extracellular matrix proteins that line and protect epithelial surfaces (20, 21). Within the mammalian digestive tract, the predominant intestinal mucin MUC2 is thought to undergo a regulated unfolding process during secretion that results in the formation of a hydrated layer that expands to line and protect epithelial cells (15, 21, 22). However, how these large mucins are packaged into secretory granules prior to secretion is unclear. In vitro studies using N-terminal portions of MUC2 suggest that pH-dependent interactions regulate the formation of supramolecular MUC2 structures (16, 23, 24). Based on these in vitro studies, one model suggests that C-terminal disulfide bonding along with Ca^2+^-dependent charge shielding may result in the compaction of MUC2 complexes into stacked rings that form bundled rod-like structures upon packaging within secretory granules (16). An alternative model suggests that MUC2 is compacted as linear polymers (24). Similar in vitro folding studies with truncated portions of the pulmonary mucin, MUC5B, have demonstrated calcium and pH-dependent N- and C-terminal interactions that are proposed to result in the formation of compacted polymers arranged around a central node to form disc-like structures (25–27). However, whether fulllength, native MUC2 or MUC5B form such structures within secretory granules in vivo and how this compaction process is orchestrated remain unknown. Also, whether multiple large proteins can be efficiently packaged and appropriately separated within the same secretory granule or whether each mucin requires its own distinct granule also remains unknown.

*Drosophila* salivary glands (SGs) are the major secretory organ in the fly and synthesize multiple highly O-glycosylated mucins that are packaged and secreted in response to developmentally-regulated hormone pulses (28–32). The size of the glands and their secretory structures, in addition to an array of the fluorescently-labeled proteins, have allowed visualization of early stages of secretory granule formation through final secretion of protein products (33–35). By following fluorescently-labeled mucin (Sgs3-GFP; (33)) in the SGs, it has been shown that secretory granule biosynthesis is dependent on the conserved ER/Golgi transmembrane protein Tango1, which forms functional contact/docking sites between the ER and Golgi, through which cargo proteins transit (36, 37). Once in the Golgi apparatus, cargo proteins are modified/glycosylated and then packaged into small secretory granules that bud from the trans-Golgi network (TGN) in a clathrin and AP-1 dependent fashion (36, 38). These small immature secretory granules then undergo homotypic fusion (36, 38–40) and maturation steps that are dependent on endosomally-derived proteins (41, 42) to form large granules that are between 3-8 μM in diameter (38–40). These large secretory granules remain in the cytoplasm awaiting a second hormone pulse, at which time the granules fuse with the plasma membrane and secrete their contents into the lumen of the SGs in an actomyosin-dependent process (34, 35). However, it is not known whether the multiple distinct mucins expressed by the SGs are targeted to unique secretory granules or packaged within the same granule. Likewise, the details of cargo compaction/segregation and the factors that control it during the secretory granule enlargement/maturation phase remain unknown.

Given our current gaps in understanding how large extracellular proteins are sorted, packaged and folded within secretory granules to ensure proper secretion in vivo, we used the *Drosophila* SG system to address these questions. Here through genetics and multimodality imaging, we find that each secretory granule contains multiple distinct mucins that undergo organized folding/restructuring events to generate distinct structures that become spatially segregated within mature secretory granules. Moreover, we define the endogenous mucins that are components of each distinct structure and further identify the temporally-regulated genes controlling pH and ion concentrations that are essential for the restructuring events occurring during the secretory granule maturation process. Our data indicate that individual mucins adopt unique structures in vivo through a regulated secretory granule maturation process controlled by calcium ions, chloride ions, pH and glycosylation. Furthermore, we show that intragranular restructuring and spatial segregation of each mucin, rather than intergranular separation of distinct mucins, is the predominant mechanism for cargo packaging and secretion within this organ.

## RESULTS

### Multiple distinct structures exist in individual secretory granules

Using fly lines that express fluorescently-labeled mucin (Sgs3-GFP) (33), we examined SGs at various stages of development based on the presence of Sgs3-GFP containing secretory granules (Fig. 1a-d). As described previously (33, 38), early 3^rd^ instar SGs that have not yet begun to synthesize secretory cargo have no GFP present (Stage 0) (Fig. 1a). As development proceeds and the first hormone pulse occurs, SGs begin to synthesize Sgs3-GFP cargo in a spatially-regulated fashion, beginning in the distal-most cells (Stage 1) (Fig. 1b) and then proceeding proximally. As shown previously, small immature secretory granules then undergo homotypic fusion to form large, mature granules that are present throughout all secretory cells of the SG (Stage 2) (Fig. 1c). Finally, in response to a second hormone pulse, secretory granules fuse with the apical plasma membrane to secrete their cargo into the lumen of the SGs (Stage 3) (Fig. 1d). We used SGs expressing Sgs3-GFP (Stage 2) (Fig.1c, e) to further investigate the details of secretory granule biogenesis and maturation via transmission electron microscopy (TEM) (Fig. 1f). Interestingly, high magnification micrographs of individual mature granules (between 3-8 μM in diameter) reveal distinct intragranular structures (Fig. 1f, g-i). For example, the majority of the granule is occupied by bundles of electron-dense filaments that are evenly spaced apart by ~28 nm (Fig. 1f, g and j). Additionally, bundles of electron-dense dots that have the same spacing seen for the filaments are also present (Fig. 1f, h and j), suggesting that they may represent cross-sections of the filament bundles. The other distinct structures observed are electron-lucent ovals/discs that are approximately 78 nm in diameter (Fig. 1f, i and k) and most often seen in close proximity to an electron-dense matrix (Fig. 1i; white arrowhead). A closer view of several high magnification TEM micrographs of the electron-lucent discs shows the presence of an electron-dense node in the center (black arrowhead in Fig. 1i). Taken together, these results demonstrate the presence of at least 3 distinct structures within mature secretory granules: filament bundles, an electron-dense matrix and electron-lucent discs. It has been suggested that aldehyde-based fixation methods could induce structural artifacts in secretory granule morphology under EM (43), However, comparison of aldehyde and cryofixed (high-pressure frozen, HPF) secretory granules demonstrates that the structures seen are independent of fixation procedures (Sup. Fig. 1a and b). Additionally, we also performed TEM on wild type (*w^1118^*) SGs that did not express the Sgs3-GFP transgene to ensure that the structures seen were not due to this transgene. As shown in Sup. Fig. 1c, the same 3 structures (filament bundles, electron-dense matrix and electron-lucent discs) were seen in the absence of Sgs3-GFP.

**Figure 1.**
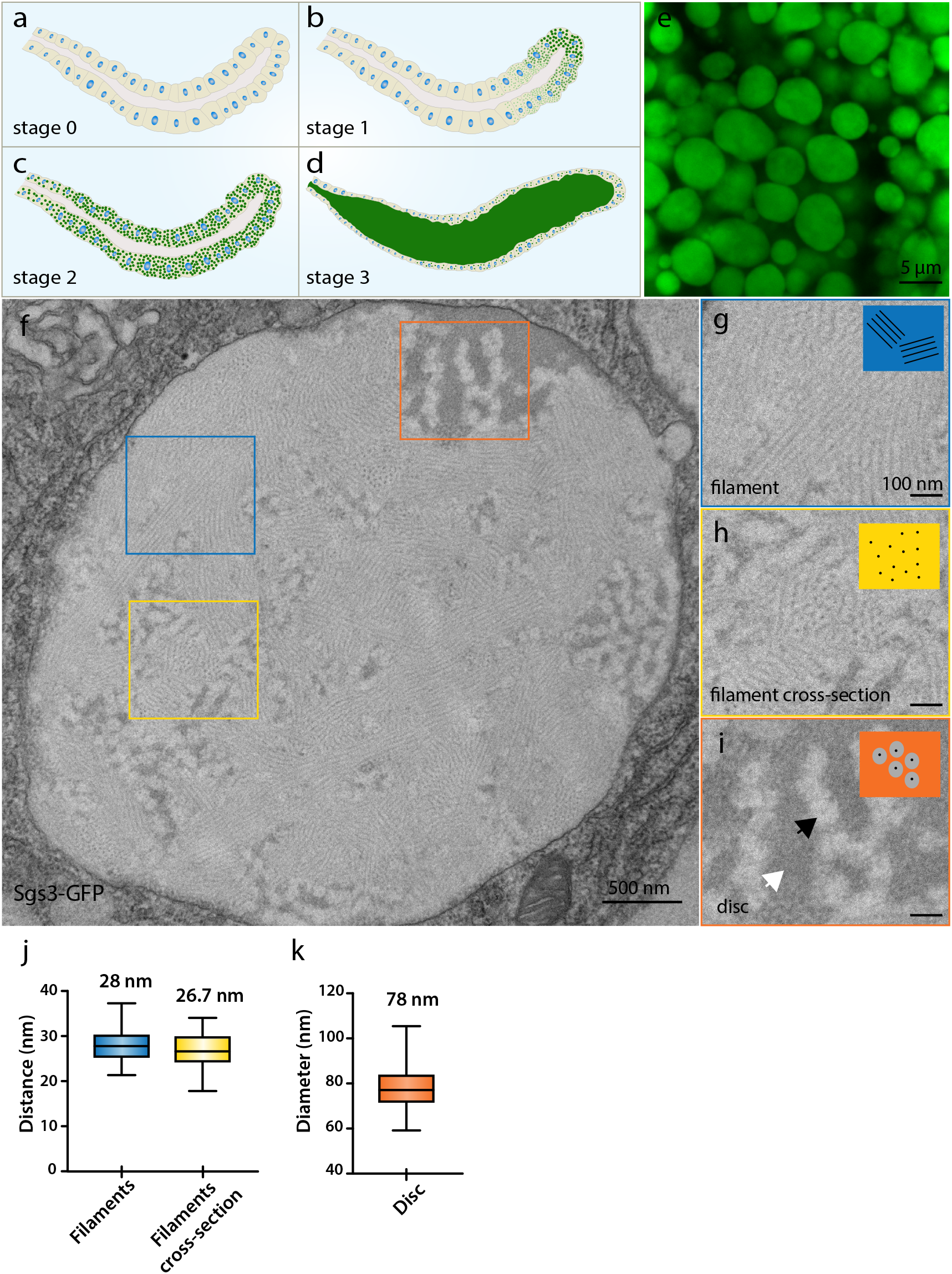
Mature secretory granules have distinct intragranular structures. Illustrations representing different stages of the third instar larval salivary glands (SGs) based on the expression pattern of Sgs3-GFP at stage 0 **(a)**, stage 1 (**b**), stage 2 **(c)** and stage 3 (**d**). Higher magnification confocal image of secretory granules at stage 2 shows their round/oval shape (**e**). Transmission electron micrographs (TEM) of a stage 2 SG show a single mature secretory granule with distinct intragranular structures **(f)**. Magnified views of the boxed regions show that the mature granule has at least three distinguishable structures: electron-dense filaments arranged in parallel bundles (**g** and **h**), electron-lucent discs (**i**; black arrowhead) and an electron-dense matrix (**i**; white arrowhead). In most of the stage 2 granules observed under TEM, the electron-lucent discs were closely associated with the electron-dense matrix (**i**; white arrowhead). Morphometric analysis of the stage 2 granules showing the inter-filament distances (**j**) and the average diameter of the discs (**k**). Scale bars, 5 μm for **e**, 500 nm for **f** and 100 nm for **g**-**i**. Representative images from three independent experiments are shown.

To better understand the spatial distribution of the electron-lucent discs (hereafter referred to as “discs”) and the electron-dense matrix, we performed Focused Ion Beam Scanning Electron Microscopy (FIB-SEM) on stage 2 SGs. We acquired a 3D volume of a single granule with an isotropic resolution of 6 nm (Fig. 2a), and used a cropped region (160 × 334 × 200, xyz) within the granule in different orientations for segmentation and 3D reconstruction (Fig. 2b and 2c-2f). Semi-automated segmentation was performed and for the purpose of illustration, the discs were pseudocolored in red and the electron-dense matrix in green (Fig. 2b’). Manual analysis of the 3D reconstruction volume reveals that the electron-dense matrix has indentations (arrows in Fig. 2g) into which the discs are embedded (Fig. 2g’-g”). The discs were found to be embedded at the peripheral surface of the electron-dense matrix when viewed at different rotational angles (Fig. 2h-j) and not within the electron-dense matrix as revealed by the orthogonal sections through the 3D volume (Fig. 2k). Therefore, our data suggest that the discs are present around the periphery of the electron-dense matrix. Taken together, our data indicate each mature secretory granule contains multiple distinct structures with defined dimensions. Moreover, and two of these structures (discs and electron-dense matrix) are associated with one another spatially to become partially segregated from the third structure (filament bundles).

**Figure 2.**
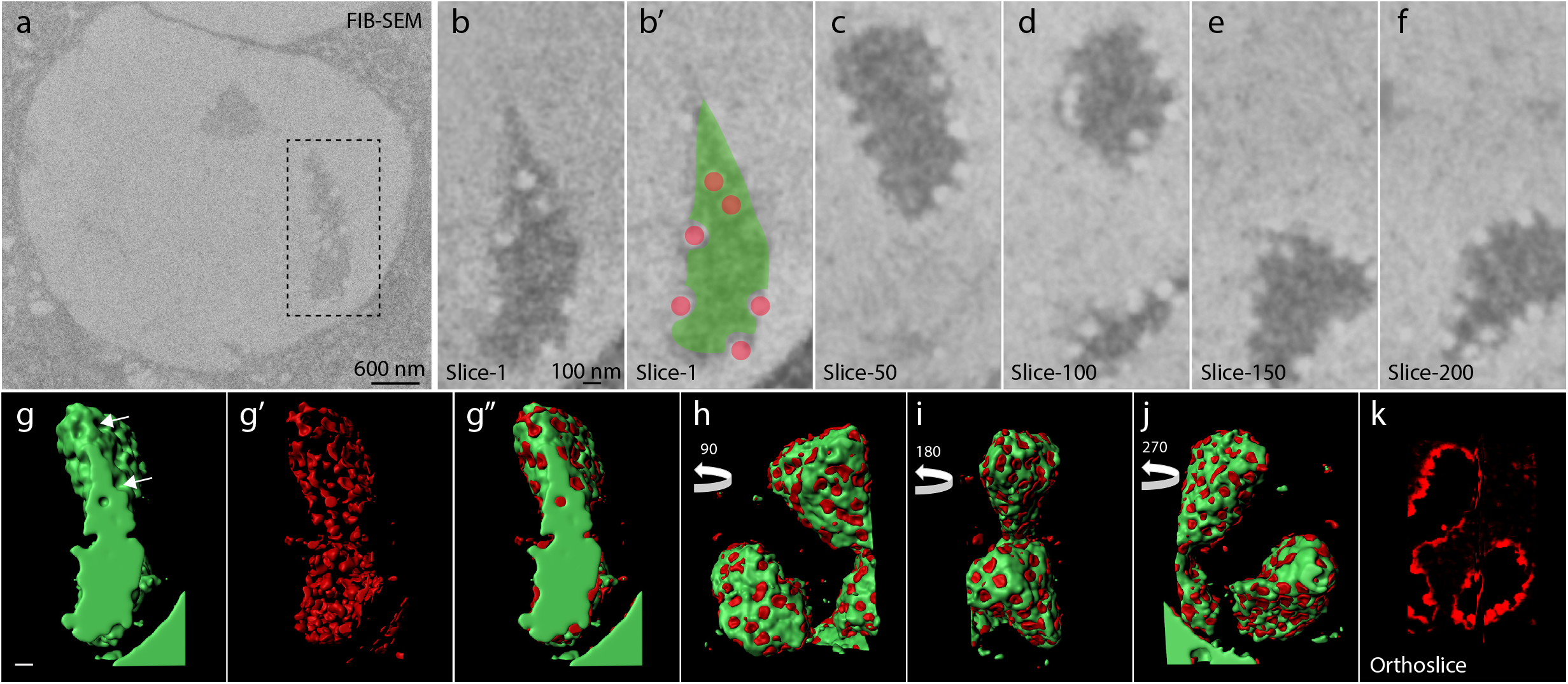
Unique associations between distinct intragranular structures. Focused ion beam scanning electron microscopy (FIB-SEM) of stage 2 secretory granules (**a**-**f**) shows slices through the region encompassing the discs and electron-dense matrix. Pseudo-colored discs (red) and electron-dense matrix (green) (**b’**) were reconstructed in 3 dimensions in **g**-**k**. White arrows in **g** highlight indented regions of the electron-dense matrix in which the discs are embedded. Images are rotated 90° (**h**), 180° (**i**) and 270° (**j**). Orthogonal slice showing only the red pseudocolored discs is shown in **k**. Scale bars, 600 nm for **a** and 100nm for **b**-**k**.

### Secretory cargo undergoes regulated restructuring during secretory granule maturation

To understand the development of the distinct structures present within mature secretory granules, we imaged secretory granules beginning at the early stages of granule biosynthesis (Stage 1) using TEM. During stage 1, the SG displays a gradient of secretory granules at various stages of maturation, with granule biogenesis beginning at the distal tip and proceeding proximally (Fig. 1b). We therefore sliced the SGs from proximal to distal (at 25 μm intervals) to capture secretory granules at various stages of development. In the initial proximal sections captured (halfway the length of SGs), we observed several small granules scattered throughout the cytoplasm ranging between 200-400 nm in diameter (Fig. 3a and Sup.Fig. 1d). These small granules often had no discernable internal structures or the beginnings of what appear to be irregularly-spaced filaments (Fig. 3a’). In subsequent sections, we observed granules ranging from 600-900 nm in diameter that contained more irregularly-spaced filaments and a few small regions with higher-electron density (Fig. 3b-b’ and Sup.Fig. 1e). Further distal sections revealed larger diameter granules (900-1200 nm) with more distinct filaments in addition to larger electron-dense regions (Fig. 3c-c’ and Sup.Fig. 1f). In sections with secretory granules ranging from 1200-1800 nm in diameter, filaments are now very distinct and arranged in evenly-spaced, parallel stacks throughout the granule along with scattered electron-lucent zones (Fig. 3d-3d’ and Sup.Fig. 1g). Additionally, very distinct and large electron-dense regions are present in these granules.

**Figure 3.**
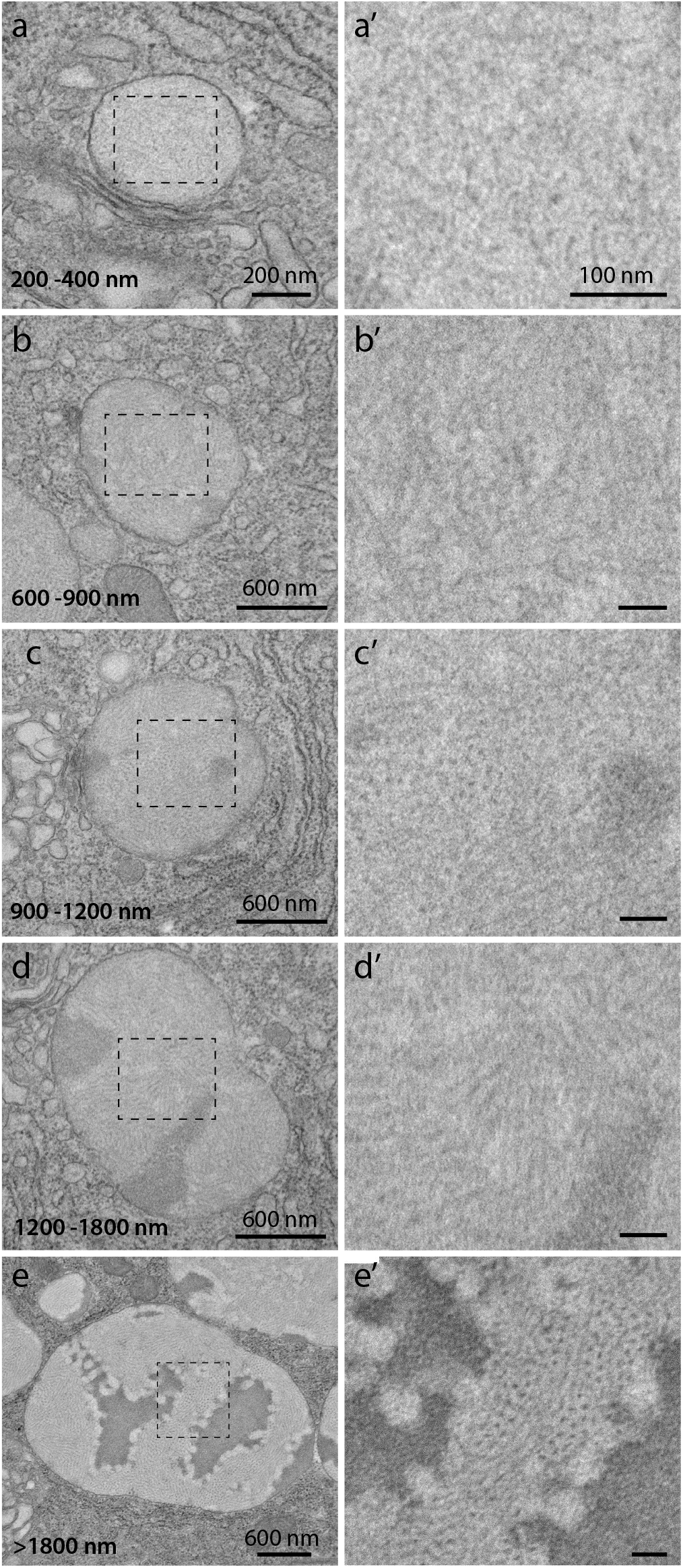
Secretory granule cargo undergoes regulated restructuring. Transmission electron micrographs (TEMs) of stage 1 and stage 2 SGs show the formation of distinct structures over time during secretory granule maturation. Early stage 1 secretory granules with diameters ranging from 200-400 nm show the first appearance of irregularly spaced electron-dense filaments (**a-a’**). As granules reach 600-900 nm in diameter (**b-b’**), the electron-dense matrix starts to appear. In the granules with diameters ranging from 900-1200 nm (**c-c’**), larger electron-dense regions and some parallel filaments become visible. Secretory granules that are 1200-1800 nm in diameter (**d-d’**) have larger areas of electron-dense regions, some electron-lucent regions and the filaments are more evenly spaced. Granules greater than 1800 nm in diameter (**e-e’**) display ordered filament bundles and distinct electron-lucent discs in close association with the electron-dense matrix, as is seen in fully mature granules. Scale bars, 200 nm for **a**, 600 nm for **b-e**, and 100 nm for **a’-e’**. Representative images from three independent experiments are shown.

We next analyzed stage 2 SGs, where most of the granules observed had diameters greater than 1800 nm (Fig. 3e-e’ and Sup. Fig. 1h). During this stage, each secretory granule has filaments arranged in evenly spaced parallel stacks. We also observed the discs that line the periphery of the distinct and large electron-dense matrix at this stage. Taken together, these results suggest that secretory cargo undergoes regulated restructuring during secretory granule maturation, with filaments forming first, followed by electron-dense aggregates and then finally discs that form along the periphery of the electron-dense regions. Moreover, our results further suggest the existence of multiple unique cargo proteins in each granule that adopt distinct structures during secretory granule maturation.

### Distinct mucins contribute to unique intragranular structures in vivo

To determine the nature of the unique intragranular structures observed in mature secretory granules in vivo, we next performed RNAi-mediated knockdown of the genes encoding the major secretory proteins/mucins known to be expressed in the SGs, *Sgs1* and *Sgs3* (28–30, 44–46). RNAi-mediated knockdown of *Sgs1 (Sgs1^TRIP.HMC02393^*) resulted in a complete loss of the disc structures typically seen in mature control granules (Fig. 4a vs. 4b). However, no discernable changes in the organization/arrangement of the bundled filaments or the electron-dense matrix were observed (Fig. 4a vs. 4b). To verify that *Sgs1* was specifically knocked down in this experiment QPCR was employed, confirming that RNAi to *Sgs1* resulted in a >95% reduction in *Sgs1* gene expression (Fig. 4c). SDS-PAGE of protein extracts from *Sgs1^TRIP.HMC02393^* SGs stained with coomassie blue demonstrated the specific loss of a high molecular weight band corresponding to Sgs1 (47) (Fig. 4d). Western blots using a lectin (*peanut agglutinin*; PNA) which is known to detect the highly O-glycosylated Sgs1 and Sgs3 proteins also showed a specific loss of the Sgs1 band upon RNAi-mediated knockdown of *Sgs1* (Fig. 4e). Finally, an antibody raised to Sgs1 (Sup. Fig. 2) also verified the specific loss of the Sgs1 band in *Sgs1^TRIP.HMC02393^* SG extracts (Fig 4f). Taken together, these results suggest that Sgs1 is a component of the discs structures seen in mature secretory granules.

**Figure 4.**
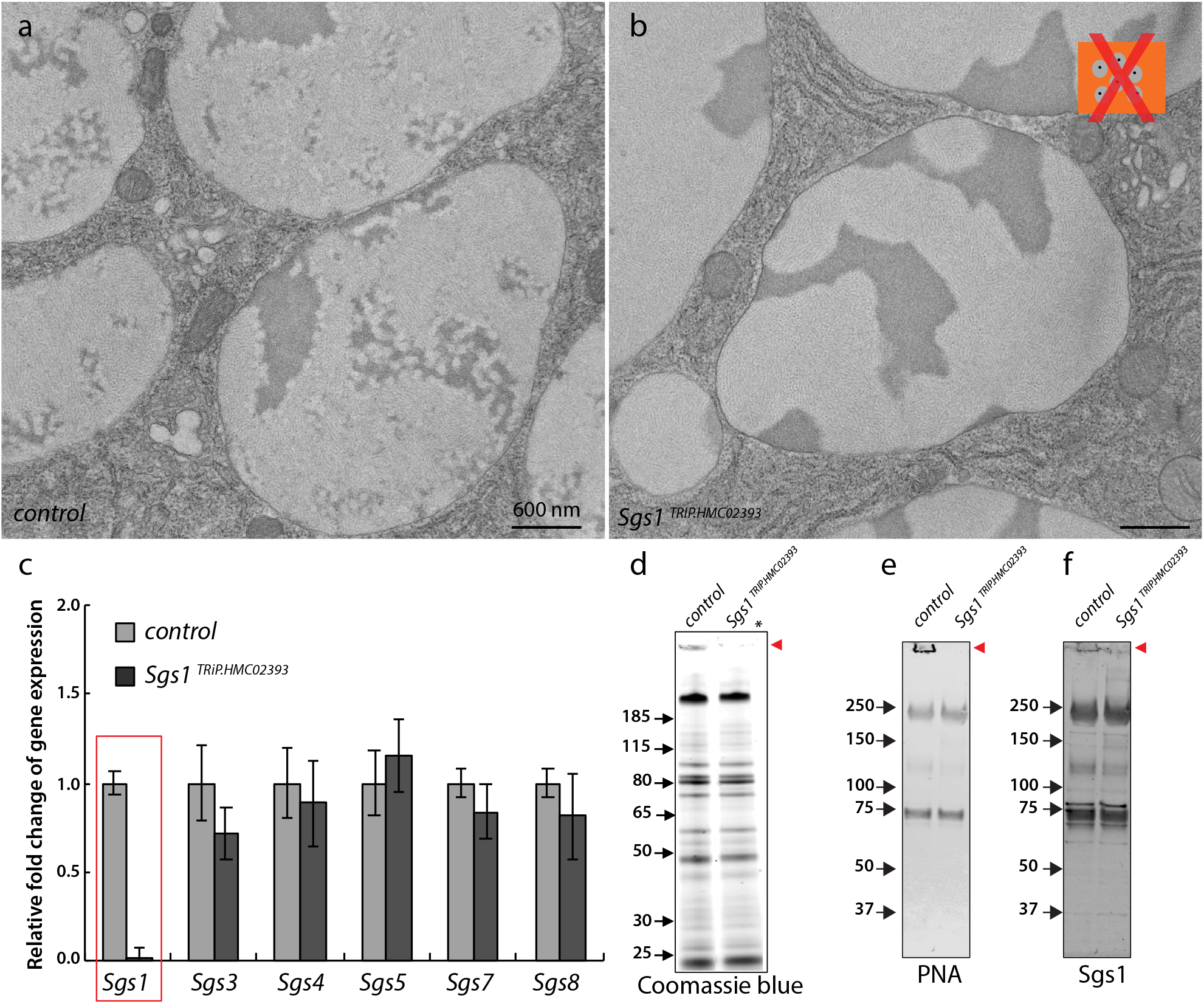
Distinct mucins form unique intragranular structures within secretory granules. TEMs on control secretory granules (**a**) were compared to those in which RNA interference (RNAi) was performed to *Sgs1* (**b**) RNAi to *Sgs1 (Sgs1^TRiP.HMC02393^*) results in the loss of the electron-lucent discs (**b**), indicating that Sgs1 is a component of the disc structures. Loss of the Sgs1 upon *Sgs1^TRiP.HMC02393^* RNAi was confirmed by QPCR (**c**) and SDS-PAGE with coomassie blue staining to show the loss of the specific Sgs1 band (**d**; red arrow). Western blots probed with the lectin peanut agglutinin (PNA), which detects the glycosylated Sgs1, shows the specific loss of the Sgs1 band (**e**; red arrow). An antibody to Sgs1 also shows loss of the specific Sgs1 band (**f**; red arrow). Scale bars, 600 nm for **a**-**b**, error bars show standard deviation in **c**. Representative images from three independent experiments are shown.

We next performed RNAi-mediated knockdown of *Sgs3 (Sgs3^TRiP.HMJ30021^*), the most abundant mucin present in the SGs. Upon performing TEM on *Sgs3^TRiP.HMJ30021^* SGs, we found complete loss of the filament bundles and the electron-dense dots (that likely represented cross-sections of the filament bundles) (Fig. 5a vs. 5b). While the filament bundles were absent, the discs and the electron-dense matrix were still present in the *Sgs3^TRiP.HMJ30021^* secretory granules. QPCR demonstrated that RNAi to *Sgs3* resulted in >95% reduction in *Sgs3* gene expression (Fig. 5c) and western blots using the PNA lectin verified the specific loss of the highly O-glycosylated Sgs3 bands (47) (Fig. 5d). Finally, westerns using an Sgs3-specific antibody ^36^ demonstrated the specific loss of Sgs3 bands in *Sgs3^TRiP.HMJ30021^* SGs (Fig.5e). Therefore, these data indicate the specificity of Sgs3 knockdown and further support the conclusion that the Sgs3 mucin is a component of the bundled filament structure within mature secretory granules. Interestingly, we also observed that the *Sgs3^TRiP.HMJ30021^* secretory granules were significantly smaller in diameter than the controls (quantitated in Fig. 5f), likely due to the specific loss of Sgs3, which is one of the most abundantly expressed proteins in the salivary gland. Attempts to immunolocalize Sgs1 and Sgs3 to structures in granules did not result in specific staining, likely because antibody binding sites are no longer accessible once these proteins have undergone restructuring and compaction within secretory granules. Taken together, our results suggest that 2 distinct mucins undergo unique packaging events within the same secretory granule to form distinct intragranular structures upon granule maturation. Additionally, given that mature control or *wt* secretory granules always contained both filaments and discs, our results support a model where distinct mucins are not sorted into unique secretory granules but rather are packaged into the same secretory granule and segregated through specific restructuring events.

**Figure 5.**
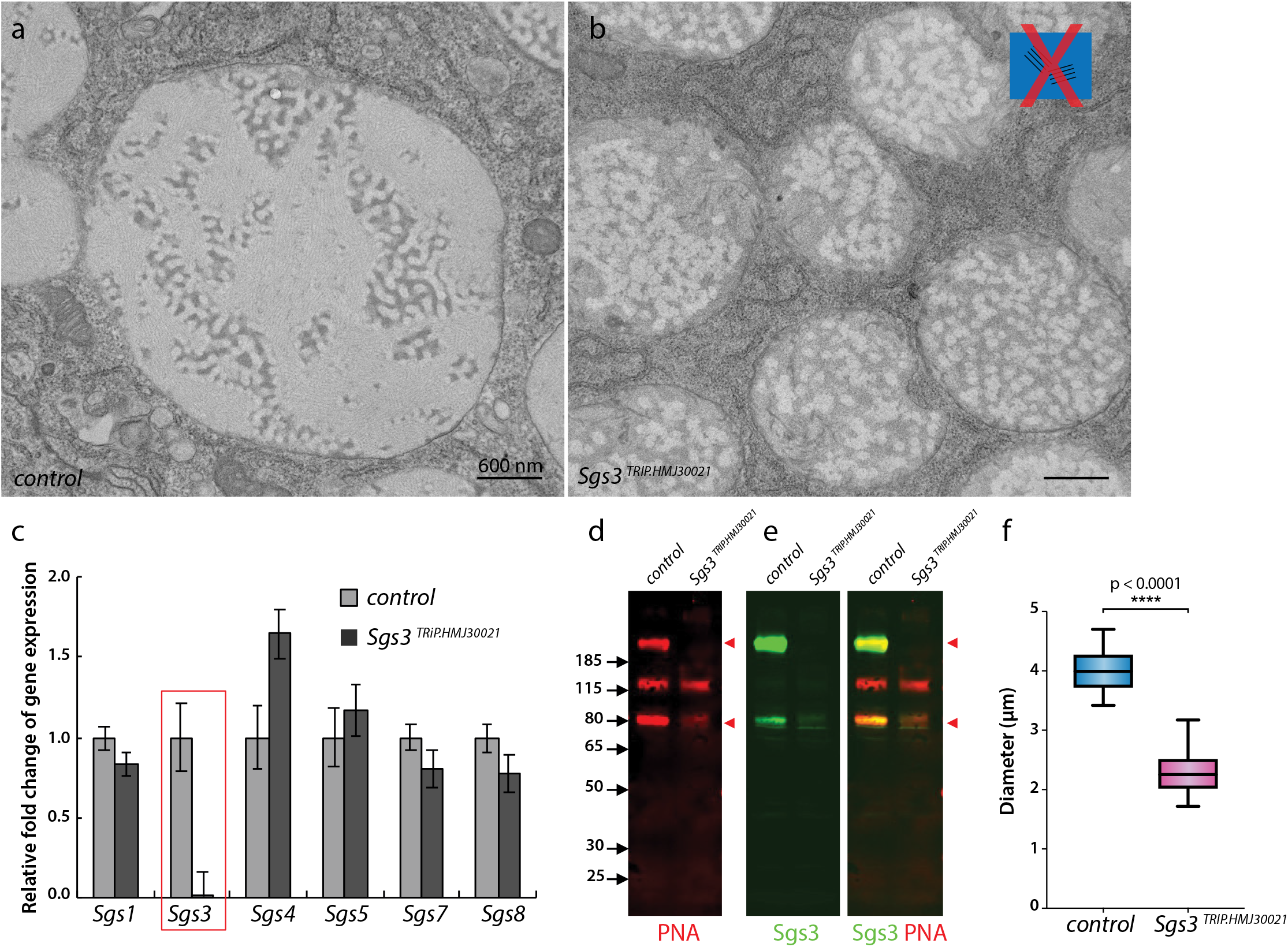
Sgs3 forms filament bundles within secretory granules. TEMs on control secretory granules (**a**) were compared to those in which RNA interference (RNAi) was performed to *Sgs3* (**b**). RNAi to *Sgs3* (*Sgs3^TRiP.HMJ30021^*) results in the loss of the filament structures that occupy most of the granule (**b**) and results in secretory granules that are much smaller in size than control. Knockdown of *Sgs3* expression was confirmed by QPCR (**c**). Loss of the Sgs3 protein upon *Sgs3^TRiP.HMJ30021^* RNAi was confirmed by western blots probed with the lectin peanut agglutinin (PNA), which showed the specific loss of the glycosylated Sgs3 bands (**d**; red arrows). Westerns probed with the Sgs3 antibody ^36^ also showed the specific loss of Sgs3 upon RNAi to *Sgs3* (**e**; red arrows). (**f**) Quantitation of the diameter of secretory granules from control and *Sgs3^TRiP.HMJ30021^* SGs is shown. ****, p<0.0001. Scale bars, 600 nm for **a**-**b**, error bars show standard deviation in **c**. Representative images from three independent experiments are shown.

### Intragranular restructuring is dependent on the v-ATPase proton pump

We next set out to interrogate the genes involved in secretory granule maturation and the restructuring of secretory cargo. To begin, we performed QPCR on SGs at various stages (stage 0, 1, 2 and 3) of secretory granule development to identify genes that are up-regulated. Previous in vitro studies have suggested that an acidic pH is required for conformational changes necessary to package large proteins, such as mucins, into more compact structures within secretory granules (16, 48). To test this hypothesis in vivo, we first examined the expression of genes that encode subunits of the v-ATPase proton pump in SGs at various stages of secretory granule development (Fig. 6a). v-ATPases are a large family of multi-subunit protons pumps, composed of two functional domains V1 and V0, that are responsible for regulation of the pH in intracellular compartments (49). Interestingly, we found that one gene encoding the v-ATPase 16kD subunit-1 (*Vha16-1*) was highly expressed during all stages of secretory granule development and peaked in expression during stage 2 (Fig. 6a). We next tested the effect of this gene on intragranular morphology by performing RNAi in SGs using 2 independent lines (VDRC# 49291 and VDRC# 104490). Each line (under the control of the *c135-Gal4* driver) showed a greater than 75% knockdown of *Vha16-1* transcripts in the SGs (Sup. Fig. 3a). Confocal imaging of stage 2 SGs dissected from *Vha16-1^VDRC49291^* or *Vha16-1^VDRC104490^* animals revealed secretory granules that were circular and swollen in appearance when compared to control granules (Fig. 6b, 6c and Sup. Fig. 3b and c). Quantitation of circularity showed statistically significant differences between control and *Vha16-1^VDRC49291^* or *Vha16-1^VDRC104490^* secretory granules (Fig. 6d). To confirm the specific role of the proton pump and pH in secretory granule morphology, we also treated SGs with the v-ATPase inhibitor Bafilomycin A1 (50), which it known to specifically inhibit the v-ATPase proton pump. Bafilomycin A1 treatment resulted in swollen and circular secretory granules (Fig. 6e vs. 6f), mimicking the phenotype seen upon knockdown of *Vha16-1*. Taken together, these genetic and pharmacologic experiments support a role for v-ATPase activity and pH in secretory granule morphology.

**Figure 6.**
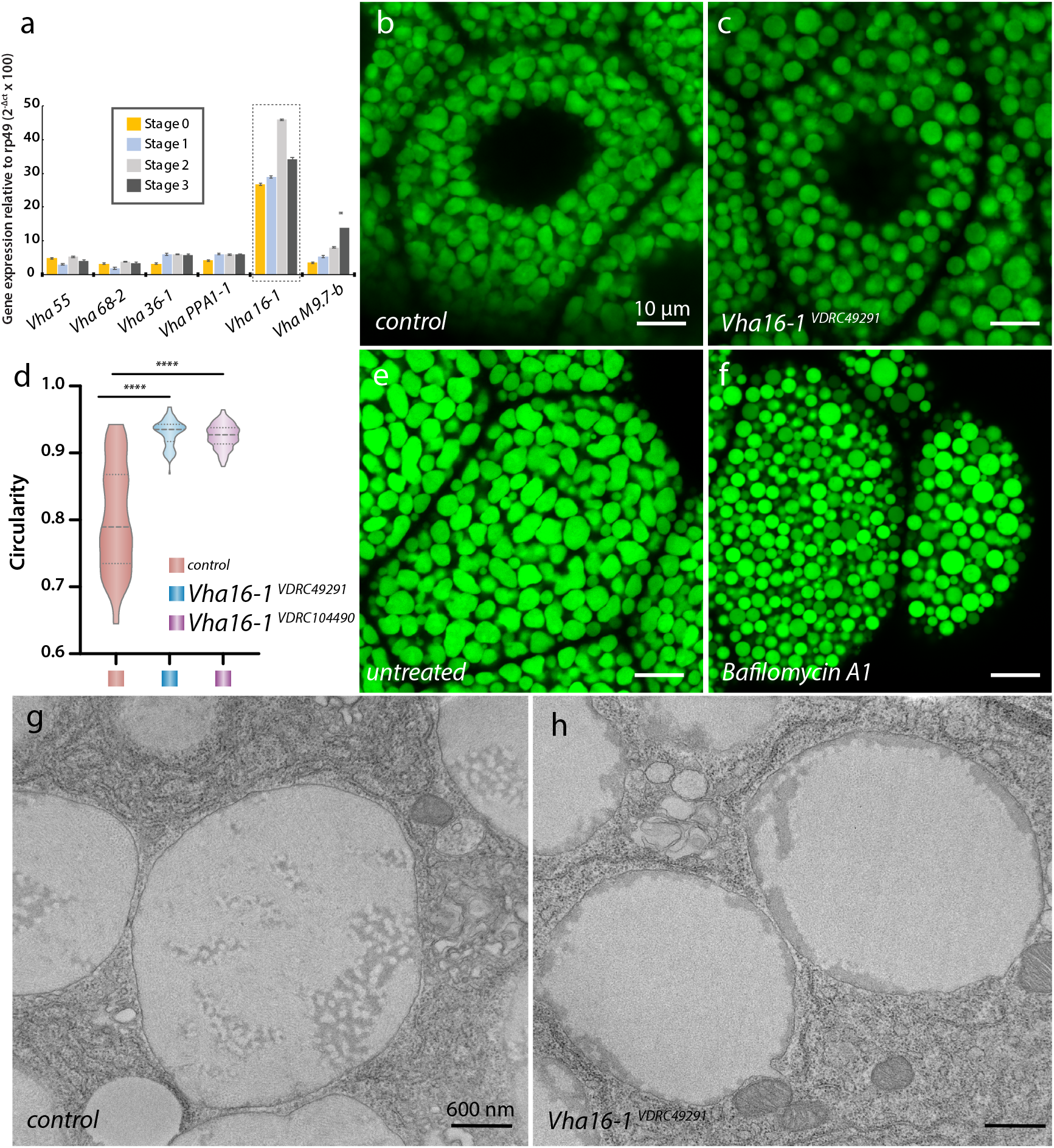
Intragranular restructuring is dependent on the v-ATPase proton pump. (**a**) QPCR analysis of the gene expression of genes encoding the subunits of the v-ATPase proton pump at each stage of SG development. Confocal images of control (**b**) and RNAi-mediated knockdown of the *Vha16.1* subunit using *Vha16.1^VDRC49291^* (**c**) are shown. Circularity measurements for two independent RNAi lines, *Vha16.1^VDRC49291^* and *Vha16.1^VDRC104490^*, are shown in (**d**). SGs untreated (**e**) or incubated with the v-ATPase inhibitor Bafilomycin A1 (**f**) recapitulate the round secretory granule phenotype seen upon knockdown of *Vha16.1*. TEMs of stage 2 control (**g**) and *Vha16.1^VDRC49291^* (**h**) SGs showing the effects of *Vha16.1* knockdown on intragranular structures, including the disruption of the Sgs3 filaments. Error bars show standard deviation in **a**, Scale bars, 10 μm for **b**-**e**, 600 nm for **g**-**h**. ****, p<0.0001. Representative images from three independent experiments are shown.

To further understand the ultrastructural changes resulting from the loss of the v-ATPase, we next performed TEM on *Vha16-1^VDRC49291^* SGs. Interestingly, we saw significant changes in intragranular morphology in the absence of *Vha16-1. Vha16-1^VDRC49291^* SGs lacked the distinct Sgs3 filament bundles, suggesting that an acidic pH is required for the formation of these structures (Fig. 6g vs. 6h). The Sgs1 discs and the electron-dense matrix were still present but were no longer aggregated within the granule, but dispersed along the periphery of the granule. These results suggest that *Vha16-1* and the changes in pH it induces are required for the appropriate folding/restructuring of Sgs3 in vivo.

### Loss of the calcium channel *fwe* disrupts secretory granule maturation

In vitro reconstitution experiments have indicated that calcium is required for proper mucin folding (16, 51, 52). To determine how calcium influences intragranular mucin morphology in vivo, we analyzed the gene *flower (fwe*), which encodes a transmembrane calcium ion channel that regulates synaptic vesicle exo- and endocytosis (53) and is upregulated in the larval SG (Fig. 7a). Confocal imaging of stage 2 SGs expressing a fluorescently tagged *fwe* genomic rescue construct (*fwe-YFP*) (53) shows that fwe localizes to secretory granule membranes (Fig. 7b). RNAi knockdown of *fwe* was performed using two independent RNAi lines (VDRC #39596 and TRiP.GL01498) (Fig. 7d and Sup. Fig. 3d-f). Confocal imaging of stage 2 *fwe^VDRC39596^* and *fwe^TRiP.GL01498^* SGs showed regions containing very large granules with low-GFP intensity (Fig. 7c vs. 7d, and Sup. Fig. 3e and 3f). TEM analysis of SGs from stage 2 *fwe^VDRC39596^* animals showed large granules that lacked the Sgs1 discs and the electron-dense matrix (Fig. 7e vs. 7f). Additionally, the organized bundled filaments typical of Sgs3 were no longer present. Instead, secretory granules were filled with very long winding filaments that spanned the entire lumen of the granule, making it look like a ball of yarn (Fig. 7e vs. 7f). Therefore, loss of *fwe* disrupts the restructuring of both Sgs1 and Sgs3, suggesting that the formation of these structures is calcium-dependent in vivo.

**Figure 7.**
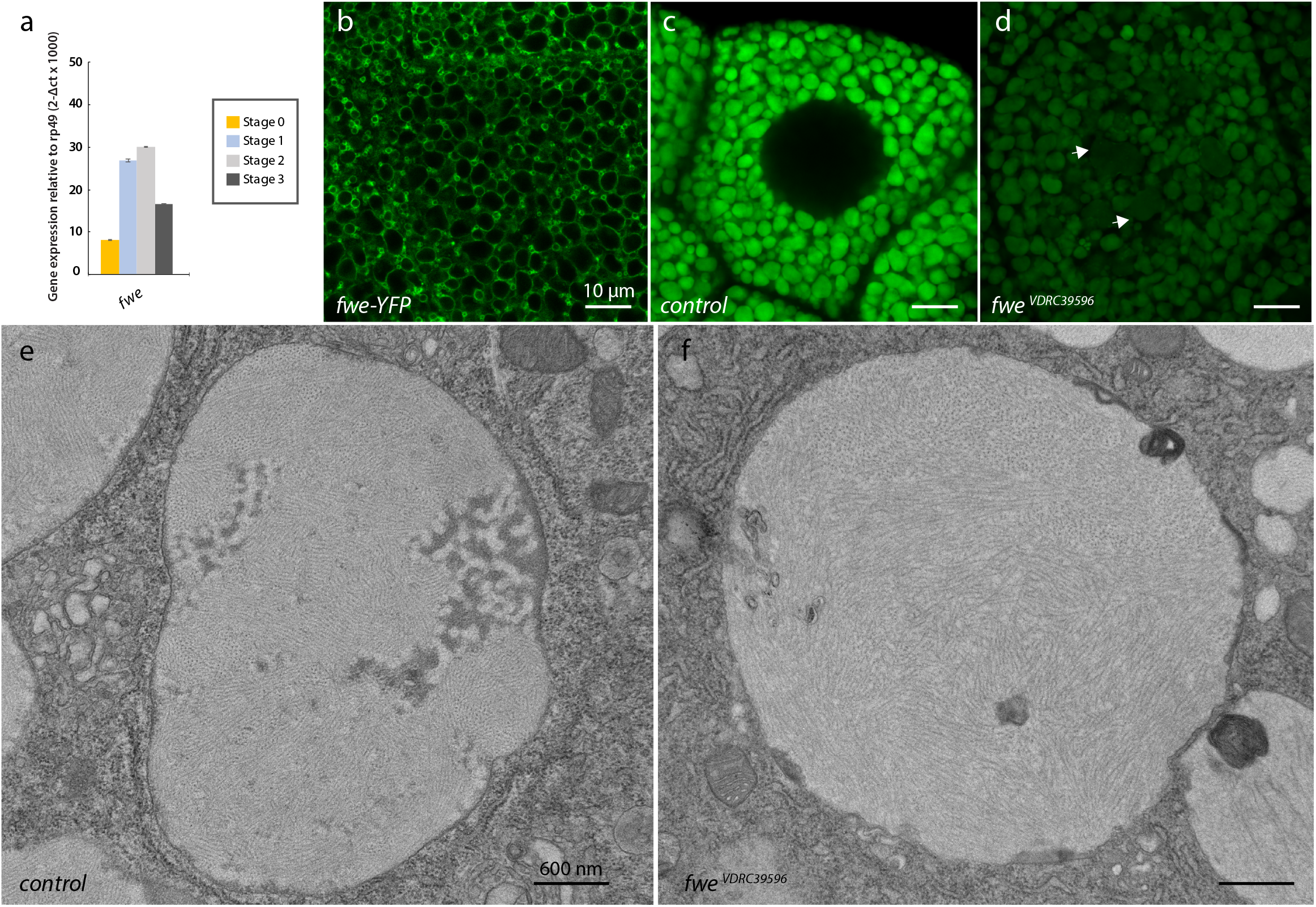
Loss of the calcium channel fwe disrupts intragranular restructuring. (**a**) QPCR analysis of the gene expression of *fwe* at each stage of SG development. The calcium channel fwe is present on secretory granule membranes as visualized in the fwe-YFP line (**b**). Confocal images of control (**c**) and RNAi-mediated knockdown of *fwe* using two independent RNAi lines, *fwe^VDRC39596^* (**d**) or *fwe^TRiP.GL01498^* (Sup. Fig. 3d-f) resulted in irregular granule morphologies. TEMs show that the loss of *fwe* (**f**) results in granules that are filled with disordered winding filaments relative to control (**e**). Error bars show standard deviation in **a**, Scale bars, 10 μm for **b**-**d**, 600 nm for **e**-**f**. Representative images from three independent experiments are shown.

### Loss of chloride ion channel proteins disrupt restructuring and intragranular morphology

Along with acidic pH, in vitro studies have suggested a role for various ions in charge shielding to promote packaging of mucin polymers (16, 51, 52). QPCR data from SGs at various stages revealed that two chloride channels (*Clic and ClC-c*) are highly expressed and upregulated during stages 1 and 2 (Fig. 8a). We therefore examined the effects of these ion channels on secretory granule morphology and cargo restructuring by performing RNAi to each. *Clic*, which encodes an intracellular chloride transporter localized to the mitochondria (54), was knocked down using 2 independent lines (VDRC #105975 and VDRC #28303). RNAi to *Clic* using either line (under the control of the *c135-Gal4* driver) gave in a greater than 75% decrease in gene expression by QPCR (Sup.Fig. 3g) and resulted in granules that were circular, swollen and much larger in diameter than control when observed using confocal microscopy (Fig. 8b vs. c and Sup. Fig. 3h and 3i). TEM examination of stage 2 *Clic^VDRC105975^* SGs revealed a notable loss of the electron-dense matrix relative to control (Fig. 8e vs. 8f). Likewise, the Sgs1 discs, which are normally associated with the electron-dense matrix, were found to be scattered throughout the granule. Some Sgs3 filament bundles could still be seen but appeared to be shorter and less distinct relative to control (Fig. 8e vs. 8f). Along with the disrupted intragranular morphology, we also saw swollen and damaged mitochondria (Sup. Fig. 4a-4d), suggesting that defects in intragranular morphology could also be the result of defective mitochondrial function.

**Figure 8.**
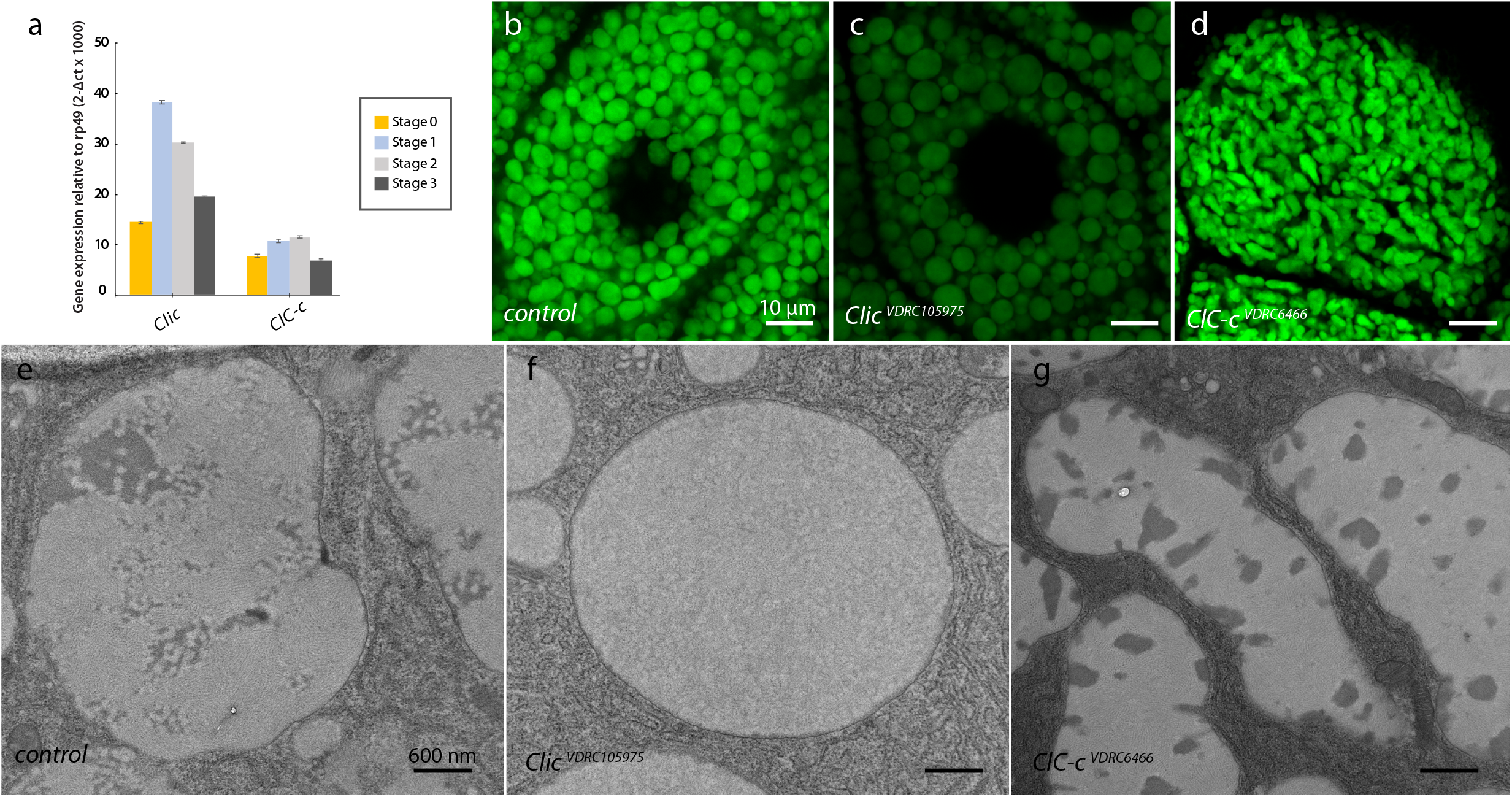
Loss of chloride ion channels alter secretory cargo restructuring. (**a**) QPCR analysis of the gene expression of *Clic* and *ClC-c* at each stage of SG development. Confocal images of control (**b**) and RNAi-mediated knockdown of *Clic* using *Clic^VDRC105975^* (**c**) or *ClC-c* using *ClC-c^VDRC6466^*(**d**) are shown. TEMs of control (**e**) and *Clic^VDRC105975^* (**f**) and *ClC-c^VDRC6466^* (**g**) granules are shown. Error bars show standard deviation in **a**, Scale bars, 10 μm for **b**-**d**, 600 nm for **e**-**g**. Representative images from three independent experiments are shown.

We next examined the role of the second-most abundantly expressed chloride channel (*ClC-c*) in intragranular morphology. *ClC-c* is a homolog of human *CLCN3(CLC-3*) which encodes a voltage-gated chloride transporter that is involved in endosomal acidification by functioning as a proton antiporter (55, 56). RNAi to *ClC-c* was performed using two independent lines (VDRC #6466 and VDRC #106844) (Sup. Fig. 3j) and resulted in granules that were grossly misshapen when examined using confocal microscopy (Fig. 8b vs. 8d and Sup. Fig. 3k vs. 3l). *ClC-c^VDRC6466^* and *ClC-c^VDRC106844^* secretory granules were oblong compared to the oval granules seen in control SGs. TEM analysis of *ClC-c^VDRC6466^* revealed major changes in intragranular structures, with the electron-dense matrix fragmented into smaller islands and scattered throughout the granule lumen (Fig. 8e vs. 8g). The Sgs1 discs were still present around the periphery of the electron-dense matrix but were much less distinct in structure. The Sgs3 filaments maintained their parallel stacked arrangement and did not show any abnormal organization. These results suggest that the loss of *ClC-c* is specifically affecting the restructuring of the electron-dense matrix and the organization of Sgs1.

### Cargo glycosylation affects specific intragranular structures and secretory granule morphology

Previous studies from our group have shown that loss of an O-glycosyltransferase (PGANT9B) results in secretory granules that are abnormal in morphology and no longer circular/oval (47). This study identified the cargo protein Sgs3 as a substrate of PGANT9B. To further examine how the loss of PGANT9B affects secretory granule morphology, we examined *pgant9* mutants (*pgant9^Δ^/Df(2R)*) by TEM. Loss of *pgant9* results in secretory granules that are abnormal in shape relative to control (Fig. 9a-d). Like th *WT* control, *pgant9^Δ^/Df(2R*) granules contained the electron-dense matrix, the Sgs1 discs and the Sgs3 filament bundles. However in *pgant9* mutants, the Sgs1 discs were less pronounced than the control (Fig. 9e vs. 9f). Additionally, the spacing between the Sgs3 filaments was significantly reduced (Fig. 9g), suggesting that O-glycosylation of Sgs3 may regulate the distance between adjacent filaments. We hypothesize that the irregular granule shapes may be due to the less mobile nature of the more tightly packed Sgs3 filaments. Indeed, examination of several TEM micrographs of *pgant9^Δ^/Df(2R*) granules suggests that the mixing of granule contents may be constrained upon homotypic fusion of granules (Fig. 9b and 9c). These observations are in line with our previous study that showed that abnormal granule morphology became evident later during granule maturation as homotypic fusion was occurring (47). Taken together, we demonstrate that multiple distinct proteins undergo ordered restructuring during secretory granule maturation that is dependent on specific genes controlling granule acidification, ion changes and cargo glycosylation. We propose a model where multiple distinct mucins are packaged into the same secretory granule, where they undergo temporally-regulated restructuring events that allow for intragranular spatial segregation (Fig. 10).

**Figure 9.**
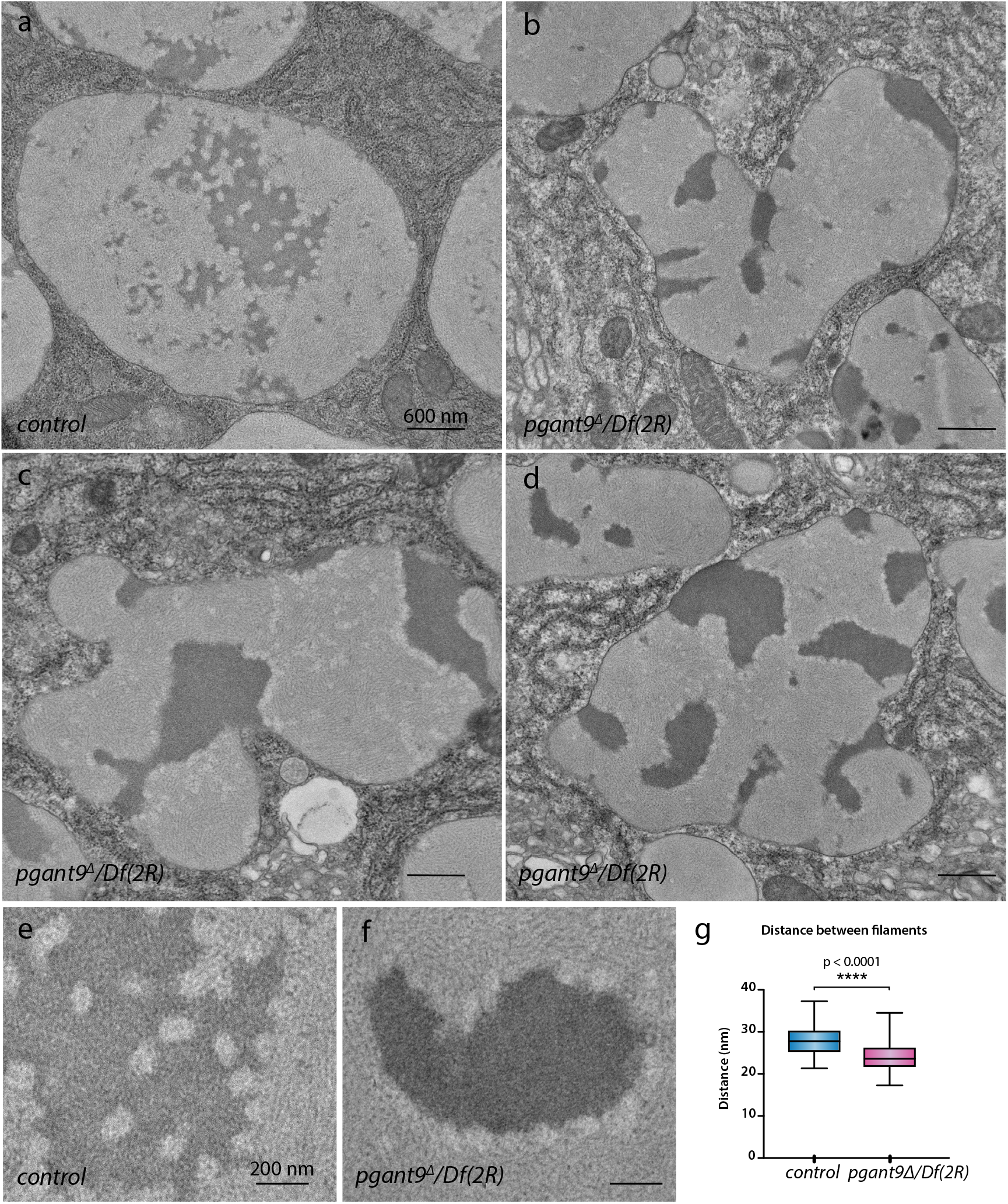
Loss of mucin-type O-glycosylation alters specific intragranular structures. TEMs of control (**a**) and *pgant9* mutant (*pgant9^Δ^/Df(2R)*) stage 2 SGs (**b-d**) are shown. Higher magnification images of *pgant9* mutants (**f**) relative to control (**e**) are shown. Morphometric analysis of Sgs3 filament spacing shows that the inter-filament distance is reduced in *pgant9* mutants when compared to control (**g**). ****, p<0.0001. Scale bars, 600 nm for **a**-**d**, 200 nm for **e,f**. Representative images from three independent experiments are shown.

**Figure 10.**
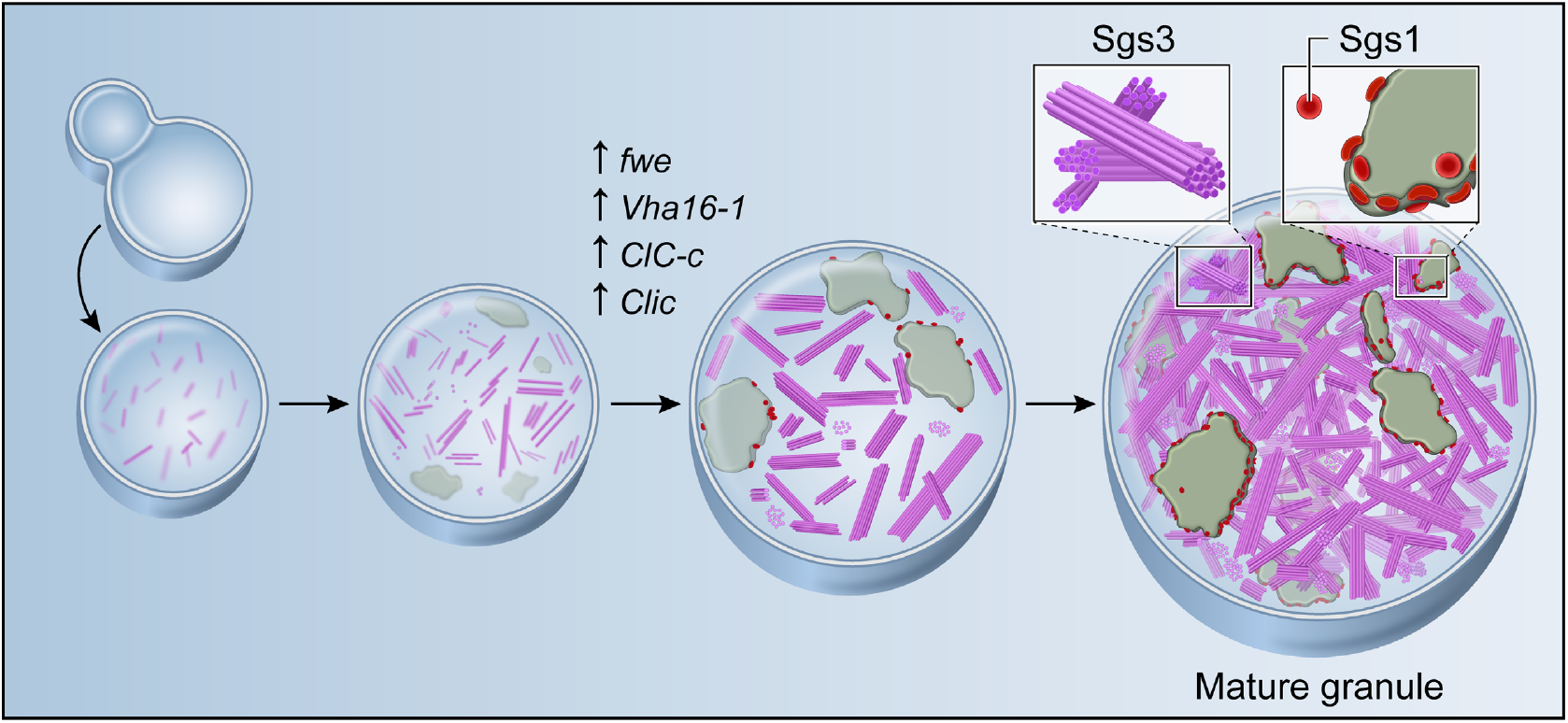
Secretory granule maturation and intragranular cargo restructuring/segregation. Secretory granule maturation involves the growth of granules by homotypic fusion along with the regulated restructuring of proteins, which is dependent on genes that control pH, calcium and chloride ions and O-glycosylation. Restructuring of the mucins Sgs1 and Sgs3 during secretory granule maturation results in the intragranular segregation of these distinct cargo proteins.

## DISCUSSION

Here we show that individual secretory granules contain multiple distinct proteins that undergo temporally-ordered restructuring during secretory granule maturation to adopt distinct intragranular structures. Moreover, we further characterize the identity of each structurally-distinct protein and some of the the factors responsible for each developing their unique structure in mature granules. Our results support a model whereby multiple cargo proteins destined to be a part of the secretate are packaged into the same secretory granule and partially segregated via orchestrated restructuring during secretory granule maturation (Fig. 10). Indeed, these results support our prior studies in the SG that show that the cargo receptor Tango1, a key protein involved in secretion, acts to form a functional docking site between the ER and Golgi that would allow protein passage between these compartments, rather than selective protein partitioning (36). The packaging of multiple proteins into a single immature granule obviates the need for specific sorting receptors for individual proteins, and instead relies on segregation via intragranular compartmentalization. Our results highlight how secretory cells under a high secretory burden may rely on intragranular rather than intergranular segregation of distinct secretory cargo.

Through a combination of genetics and ultrastructural analysis of secretory granules at different stages of development, we identify two of the unique structures as being composed of the major secretory mucins, Sgs1 and Sgs3 (57). Previous studies using a Sgs3-GFP reporter have shown that immature granules grow in size and mature by undergoing homotypic fusion (36, 38–40). Maturation of immature secretory granules by homotypic fusion has also been reported in endocrine and neuroendocrine cells (58, 59). When examining nascent granules, at different developmental stages, we did not find any containing exclusively Sgs1 or Sgs3 structures. Rather, the intragranular structures formed by these two mucins became evident gradually over time as the granules matured, suggesting that multiple mucins are packaged into a single secretory granule and that each undergoes regulated restructuring during secretory granule maturation to adopt a distinct intragranular structure. Interestingly, each mucin adopted a structure reminiscent of structures seen in vitro reconstitution experiments performed with segments of mammalian mucins. Sgs1 adopted a disc-like structure, similar to that seen upon reconstitution of a portion of MUC5B in vitro, where head-to-head N-terminal stacking is proposed to account for this structure (25–27). Likewise, Sgs3 is restructured into parallel filament bundles, similar to those seen upon in vitro reconstitution of portions of MUC2 (16, 24). The importance of the packaging of MUC2 is evident when one considers that this mucin, like many mucins that line and protect epithelia, must undergo regulated expansion into a net-like structure that lines the intestinal environment, conferring protection from microbial and mechanical damage (21). Similar to mammalian mucins, *Drosophila* mucins also form a lining that mediates attachment to a substrate to enable metamorphosis, suggesting that their packaging and expansion upon secretion must also be similarly regulated.

We have further identified the factors that mediate the ordered restructuring of the *Drosophila* mucins during secretory granule maturation in vivo. The intermediate steps of secretory granule formation/granule maturation after export from the trans-Golgi are poorly understood for mammalian mucins, although roles for calcium ions and pH have been shown to be important for this process in vitro (48, 52). Indeed, we find that two genes controlling pH and calcium concentration (*Vha16-1* and *fwe*, respectively) are differentially regulated during secretory granule maturation and are essential for mucin restructuring in vivo. Loss of *Vha16-1*, which encodes a subunit of the v-ATPase proton pump, disrupted the restructuring of Sgs3 and the association of Sgs1 discs with the electron-dense matrix. Similar disruptions were seen with a pharmacologic inhibitor of the v-ATPase, leading to grossly abnormal secretory granules and highlighting a crucial role for the proton pump and pH changes in the restructuring/segregation of cargo proteins in vivo. Loss of *fwe*, which encodes a calcium channel present on the granular membrane, resulted in the complete disruption of all intragranular structures, highlighting the crucial importance of calcium regulation in the formation of these structures and the spatial segregation of cargo. Biophysical and single-particle EM analysis proposed that the MUC2 mucin is packaged as concatenated polygonal rings, with the mucin-domains projecting perpendicular from the base of the rings as parallel rods in a calcium and pH-dependent manner within secretory granules (16). An alternative model suggests that MUC2 complexes exist as linear polymers (24) that are also calcium and pH-dependent. Previous studies utilizing single-particle EM analysis also identified calcium-mediated cross-links in the VWD-3 domain of MUC5B that could mediate stacking and condensation of the mucin into a round disc-like structure around a central node (25–27). Taken together, our in vivo studies further support a crucial role for pH and calcium in intragranular packaging of endogenous mucins and are well-aligned with previous reconstitution/folding studies that have highlighted a role for calcium and pH in the restructuring of mucins in vitro (16, 25–27).

In addition to genes controlling pH and calcium, we found that two chloride channel proteins (Clic and ClC-c) were also upregulated during secretory granule maturation and had effects on cargo morphology. The CLIC (*Cl^−^* intracellular channel) proteins are a highly conserved family of dimorphic proteins that exist in soluble and membrane forms (60). CLICs are multifunctional proteins that have been reported to play roles in apoptosis (61), membrane trafficking (62), tubulogenesis (63) and cell differentiation (64), and display differential localization (65) on nuclear membranes (66), the ER (67), mitochondria (54) and secretory vesicles of hippocampal neurons (68). *Drosophila* encodes a single homolog of CLIC (69) and knockdown of *Clic* in SGs resulted in swollen secretory granules with disrupted intragranular morphology and damaged mitochondria. The round and swollen granules with disorganized intragranular morphology seen upon knockdown of *Clic* were reminiscent of the phenotype seen upon the loss of *Vha16-1*. Since *Cl^−^* is considered to provide an electrical shunt for the acidification of most organelles of the secretory and endocytic pathway, it is conceivable that Clic might be affecting intragranular morphology by indirectly regulating pH. The other chloride channel upregulated during secretory granule maturation, ClC-c, is a homolog of the human CLC-3, which functions as an electrogenic 2Cl^−^/H^+^ exchanger (55), assists in acidification of endosomes by supplying countercurrents for the vesicular H^+^-ATPase (70), and has also been proposed to be involved in vesicular *Cl^−^* accumulation and ion homeostasis (71). In our *ClC-c* knockdown, we find a severe defect in the shape of the granule, fragmentation of the electron-dense matrix and mislocalization of Sgs1 discs. Loss of *ClC-c* may impair *Cl^−^* accumulation, leading to alterations in pH of the granule during the maturation steps. However, the loss of *ClC-c* results in a distinct phenotype from those seen upon loss of *Vha16-1* or *Clic*, suggesting a unique role for this ion channel.

Finally, we demonstrate that O-glycosylation, a protein modification abundant on mucins (72, 73), influences the structures of the mucins, thereby influencing secretory granule morphology. Previous studies from our group demonstrated that loss of one glycosyltransferase (PGANT9) that glycosylates Sgs3 resulted in secretory granules that displayed irregular morphologies (47). In the present study, ultrastructural analysis demonstrated that the loss of *pgant9* results in Sgs3 filaments that are closer together. Given that the abnormal secretory granule morphology is most evident once granules begin to undergo homotypic fusion and mature, we hypothesize that the irregular granule shapes may be due to the less mobile nature of the more tightly packed Sgs3 filaments.

In summary, we demonstrate that multiple proteins undergo temporally-regulated restructuring during secretory granule maturation to form unique structures and that these restructuring events are both pH and ion-dependent. We propose a new model for regulated intragranular cargo segregation to ensure efficient storage and secretion of large extracellular proteins such as mucins that function in the protection of epithelial surfaces. This cargo segregation in the absence of membranes may result in a type of phase separation that is occurring within secretory granules containing multiple distinct cargos. Future studies will examine how mucin packaging and the factors that influence it affect subsequent mucin unfolding/expansion during secretion.

## METHODS

### Fly strains

All fly strains and crosses were raised and maintained on MM fly media (KD Medical, Inc.), the ambient conditions were set to a 12:12 hours of dark and light cycle at 25°C. The following stocks were obtained from the Bloomington *Drosophila* stock center, *c135-Gal4* (BL6978, *w^1118^;P{w^+mW.hs^=GawB}c135), Sgs3-GFP* (BL5885, *w[*]; P{w[+mC]=Sgs3-GFP}3), Sgs1-RNAi* (BL51421, *y[1] v[1]; P{y[+t7.7] v[+t1.8]=TRiP.HMC02393}attP2), Sgs3-RNAi* (BL62944, *y[1] v[1]; P{y[+t7.7] v[+t1.8]=TRiP.HMJ30021}attP40), fwe-YFP* (BL51615, *w[*]; P{w[+mC]=fwe-8.2g-YFP-GFP-RFP}22-1-2), fwe-RNAi* (BL43157, *y[1] v[1]; P{y[+t7.7] v[+t1.8]=TRiP.GL01498}attP40), pgant9* deficiency line (BL7546, *Df(2R)Exel6064), pgant9^Δ^ (47*)). RNAi stocks expressing hairpin double stranded RNA was obtained from the Vienna *Drosophila* RNAi Center (VDRC), stocks used are VDRC#60000 (*w^1118^*), VDRC#49291 and VDRC#104490 for *CG3161 (Vha16-1*), VDRC#6466 and VDRC#106844 for *CG5284 (ClC-c*), VDRC#105975 and VDRC28303 for *CG10997 (Clic*) and VDRC#39596 for *CG6151 (fwe*). The *c135-Gal4* driver stock was recombined with *Sgs3-GFP* to generate *c135-Gal4, Sgs3-GFP* to visualize granules and induce RNAi in SGs. *c135-Gal4, Sgs3-GFP* crossed to VDRC#60000 (*w^1118^*) was used as control for the RNAi experiments.

### Transmission electron microscopy

SGs were dissected in ice-cold Schneider’s medium and fixed in 4% formaldehyde/ 2% glutaraldehyde/0.1M sodium cacodylate buffer (pH 7.4) overnight at 4 °C. After overnight fixation, salivary glands were postfixed in 1% osmium tetroxide for 1 hour on ice and overnight in 1% uranyl acetate at 4°C. The following day SGs were dehydrated in graded ethanol series followed by propylene oxide and gradually infiltrated with an increasing ratio of EMbed 812 resin (EMS, #14120) to propylene oxide with final infiltration in 100% resin. Next, SGs were embedded in Beem capsules (EMS, #70000, Size 00) and polymerized in the oven at 60C for 48 hours. Ultrathin sections (70nm) were cut on ultramicrotome (Leica EM UT7) and Digital micrographs were acquired on JOEL JEM 1200 EXII operating at 80Kv and equipped with AMT XR-60 digital camera. For High-pressure freezing and free substitution, SGs were dissected in Schneider’s medium and transferred to a Brass disc (type-A planchet, cavity depth 0.2 mm). Excess of Schneider’s medium was carefully removed and immediately 10μl of 20% BSA was pipetted and uniformly distributed. After inspecting sample orientation, another Brass disc (Type-B planchet) was placed on top with flat surface down to seal the assembly. The assembled specimen chamber was frozen using a high-pressure freezing system (Leica EM ICE). The frozen sample were transferred to cryovials in liquid nitrogen vapor and transferred to a pre-cooled (−90°C) freeze substitution unit (Leica EM AFS). Freeze substitution was performed using a mixture of 0.12% aqueous Uranyl acetate in anhydrous acetone using the following program: −90°C for 44 hrs followed by slow warming from −90°C to −47°C (15°C/hr) and at −47°C freeze-substitution solution was removed and samples were washed 3 times for 10 mins with acetone. Resin infiltration was performed by incubating samples in increasing concentrations of Lowicryl HM20 resin (25%, 50%, 75%) in acetone with final 3 incubations with 100% resin lasting for 48 hrs. The samples were gradually warmed from −47°C to 24°C (5°C/hr) and polymerized under UV over a period of 48 hrs. Ultrathin sections (70nm) were cut on ultramicrotome (Leica EM UT7) and post-stained with 2% of uranyl acetate for 10 mins and lead citrate for 2 mins. Digital micrographs were acquired on JOEL JEM 1200 EXII operating at 80 kV and equipped with AMT XR-60 digital camera.

### FIB-SEM imaging and segmentation

FIB-SEM imaging was performed as described in (74), the acquired dataset of a single granule was cropped (160 × 334 × 200, xyz) to include electron-dense matrix and discs and used further for segmentation. Image processing was peformed using Fiji (NIH, ImageJ.net), for the initial image processing, variable contrast was adjusted using the Enhance Local Contrast (CLAHE) plugin and a Gaussian blur 3D (σ =2) was applied to the image stack. Feature segmentation (electron-dense matrix and discs) was performed using the Trainable Weka segmentation plugin (version 3.2.34), three classifiers were trained to include electron-dense matrix, discs and the filaments using the FastRandomForest classifier. The resulting probability maps after training were manually inspected and the classifiers were retrained to minimize feature overlap. After refinement of the model, probability maps were exported as 32-bit hyperstack. 3D visualization, pseudo coloring and volume rendering of probability maps of the individual features was performed using Imaris (version 9.3.0, Bitplane.com).

### Antibody preparation and validation

Polyclonal Sgs1 was raised in rabbits using the peptide RTTRRRPTTPKTPD (Genscript) and was affinity purified (GenScript). To test the Sgs1 antibody, *Sgs1* cDNA 1-657bp was cloned into the EcoRI and NotI sites of pIB-V5/His vector, fused with V5 tag. Then the S2R+ cells were transfected with plasmids using Effectene transfection reagent (Qiagen) according to the manufacturer’s instructions. After 3-4 days, the cells were collected and lysed with RIPA buffer (Sigma) supplemented with 1X Halt protease inhibitor (Thermo Fisher Scientific). The proteins were electrophoresed in the 4-12% Bis-Tris gel (Invitrogen) and proteins were transferred to nitrocellulose membranes (Invitrogen, 0.45 μM pore size). After blocking with Odyssey Blocking Buffer (Li-COR), the membranes were incubated with anti-Sgs1 antibody (dilution 1:500) and anti-V5 antibody (dilution 1:1000, Thermo Fisher Scientific) at 4°C overnight. After washing, the membranes were incubated with IRDye 680LT conjugated anti-mouse IgG (1:5000, Li-COR) and 800CW conjugated anti-rabbit antibody (dilution 1:5000, LI-COR). Membranes were washed with PBST (0.1% Tween-20), rinsed in PBS, and scanned using a Li-COR Odyssey Infrared Imaging System.

### Western blotting

SGs were dissected from third instar larva and squished in 50 μl RIPA buffer (Sigma) supplemented with 1X Halt Protease Inhibitor (Thermo Scientific). Protein extracts were loaded onto NuPAGE 4–12% Bis-Tris gel (Invitrogen). For Coomassie blue staining, the gel was rinsed with ultrapure water three times and stained with SimplyBlue Safestain (Thermo Scientific). For western blotting, proteins were transferred from gel onto nitrocellulose membranes (Invitrogen, 0.45 μM pore size). The membranes were blocked with Odyssey Blocking Buffer (Li-COR) and incubated with IRDye 800CW-conjugated PNA (1:5000) and the anti-Sgs1 antibody (1:2000) or anti-Sgs3 antibody (1:2000) overnight at 4 °C. After washing with PBS containing 0.1% Tween-20 (PBST), the membrane was incubated with IRDye 680LT-conjugated antirabbit IgG (1:10000, Li-COR) for 1hr at room temperature then washed with PBST and PBS, then scanned by a Li-COR Odyssey Infrared Imaging System.

### Real-time PCR

SGs were dissected in Schneider’s medium and 10 pairs of SGs were pooled in DNA/RNA shield. RNA was extracted using the Direct-zol RNA Microprep kit (ZymoResearch) following the manufactures instructions and cDNA was synthesized from 1μg of total RNA using iScript cDNA Synthesis Kit (Bio-Rad). QPCR was performed on a CFX96 real-time PCR thermocycler (Bio-Rad) using the SYBR-Green PCR Master Mix (Bio-Rad). Primers used for QPCR are listed in Supplemental.Table 1.

### Morphometric analysis

ImageJ was used to perform morphometric analysis on the acquired TEM micrographs. Briefly, micrographs were calibrated, and the inter-filaments distance and the disc diameter were measured using the straight line and oval selection tools. A total of 100 measurements were performed across 10 micrographs and GraphPad Prism was used to perform statistical analysis. Student *t*-test was used to compare the inter-filaments distance between control and *pgant9* mutants and significant differences are reported for p-values <0.01. Circularity measurements of the secretory granules was performed as previously described (47)

### SGs imaging, Lysotracker and v-ATPase inhibitor treatment

SGs were imaged as previously described in (35). SGs were staged according to *Sgs3-GFP (33*) reporter and non-secreting glands were dissected in Schneider’s medium (Thermo Scientific) and transferred to glass-bottom culture dishes (MatTek). A polycarbonate membrane filter (Sterlitech) was gently placed on the top of the glands and the samples were imaged on Nikon A1R+ confocal microscope equipped with a Plan-Apochromat IR 60x/1.27 NA water immersion objective (For LysoTracker or v-ATPase inhibitor treatment, SGs were dissected in Schneider’s medium and incubated with Bafilomycin A1 (500 nM) or LysoTracker (1μM) in Schneider’s medium for 1 hour at room temperature. After incubation, SGs were rinsed with Schneider’s medium and transferred to glass-bottom culture dishes (MatTek) and imaged as described above. Acquired images were processed in Adobe Photoshop CC and compiled into panel using Adobe Illustrator CC 2019.

## Data availability

Data supporting the findings of this manuscript are available from the corresponding author upon reasonable request.

## Acknowledgements

We would like to thank our colleagues for many helpful discussions. We also thank the Bloomington Stock Center, the Developmental Studies Hybridoma Bank and the Vienna *Drosophila* RNAi Center for providing fly stocks, antibodies and other reagents. This research was supported by the Intramural Research Program of the NIDCR at the National Institutes of Health (Z01-DE-000713 to K.G.T.H.) and the NIDCR Imaging Core (ZIC DE000750-01).

## Author contributions

Z.A.S., L. Z., D. T. T. and K.G.T.H. designed and planned the research. Z.A.S. and L. Z. performed the experiments. Z. A.S., L. Z., D.T.T, C.K.E.B, and K.G.T.H. analyzed and discussed the data. Z.A.S. and K.G.T.H. wrote the paper.

## SUPPLEMENTARY FIGURE LEGENDS

**Sup. Fig.1:** Distinct intragranular structures observed in the mature secretory granules are unaffected by the fixation method. Both chemical fixation (**a**) and High-Pressure freezing and freeze-substitution (**b**) preserve and maintain the intragranular morphology and the observed structures are not due to any fixation induced artifacts or by the presence of *Sgs3-GFP* transgene (**c**). The distinct structures within the secretory granule undergo regulated restructuring and this gradual progression of cargo segregation is observed in secretory granules at different stages of development (**d-h**).

**Sup. Fig.2**: Validation of the Sgs1 antibody. The protein domain organization of the full length Sgs1 protein and the truncated version tagged with V5 tag (**a**). The truncated version (Sgs1p) was transfected into S2R+ cells and western blots probed with Sgs1 and V5 antibody specifically detects Sgs1p (**b**; red arrow).

**Sup. Fig.3:** QPCR confirmation of the knockdown efficiency and the Confocal images of *Vha16-1* (**a-c**)*, fwe* (**d-f**)*, Clic* (**g-i**) and *ClC-c* (**j-l**). Two independent RNAi lines were used for each gene which showed more than 75% decrease in gene expression. Ct values are normalized to *rp49* and bar graph plotted as gene expression relative to *rp49*, error bars show standard deviation. Representative images from three independent experiments are shown.

**Sup. Fig.4:** TEM micrographs comparing the control (**a**) and *Clic^VDRC105975^* RNAi (**b**) showing swollen and damaged mitochondria (black arrows) in the *Clic* deficient salivary glands, in addition to disrupted intragranular morphology. High magnification image of individual mitochondrion highlighting the extent of mitochondrial damage observed upon loss of *Clic* (**c vs. d**).

