## Supplementary figures and images for "Orchestrated Restructuring Events During Secretory Granule Maturation Mediate Intragranular Cargo Segregation"

### Sup. Figs. 1-4

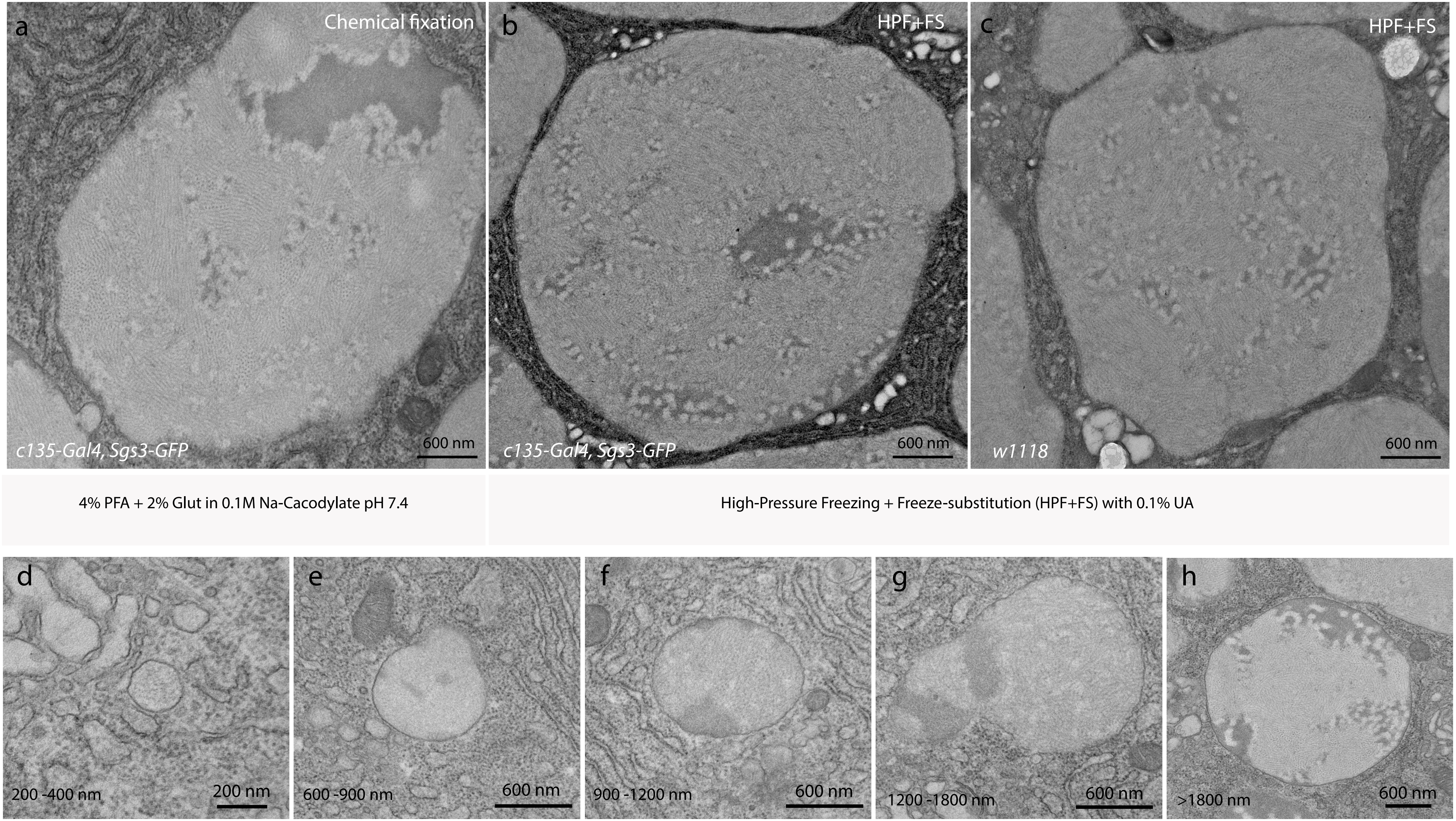

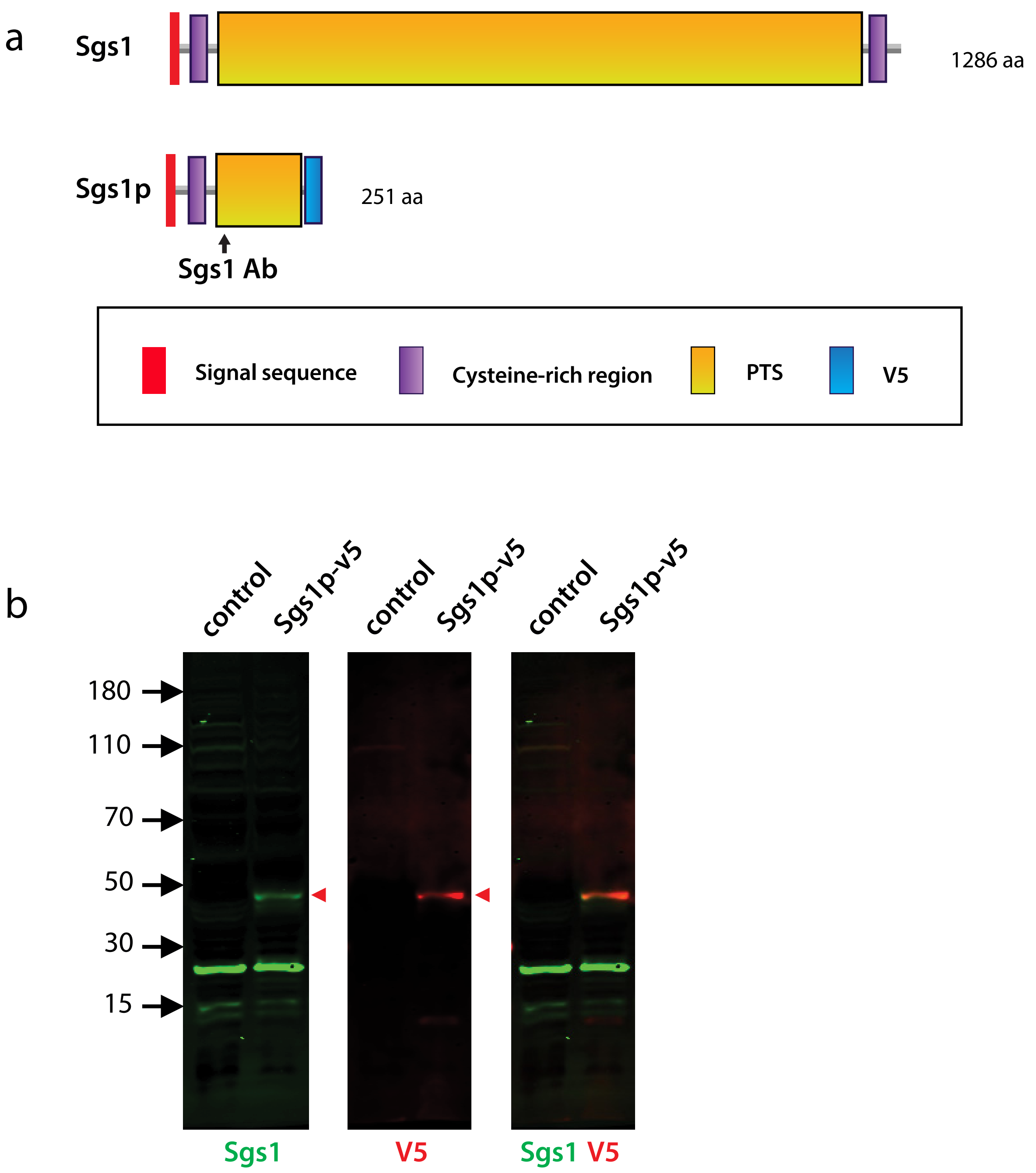

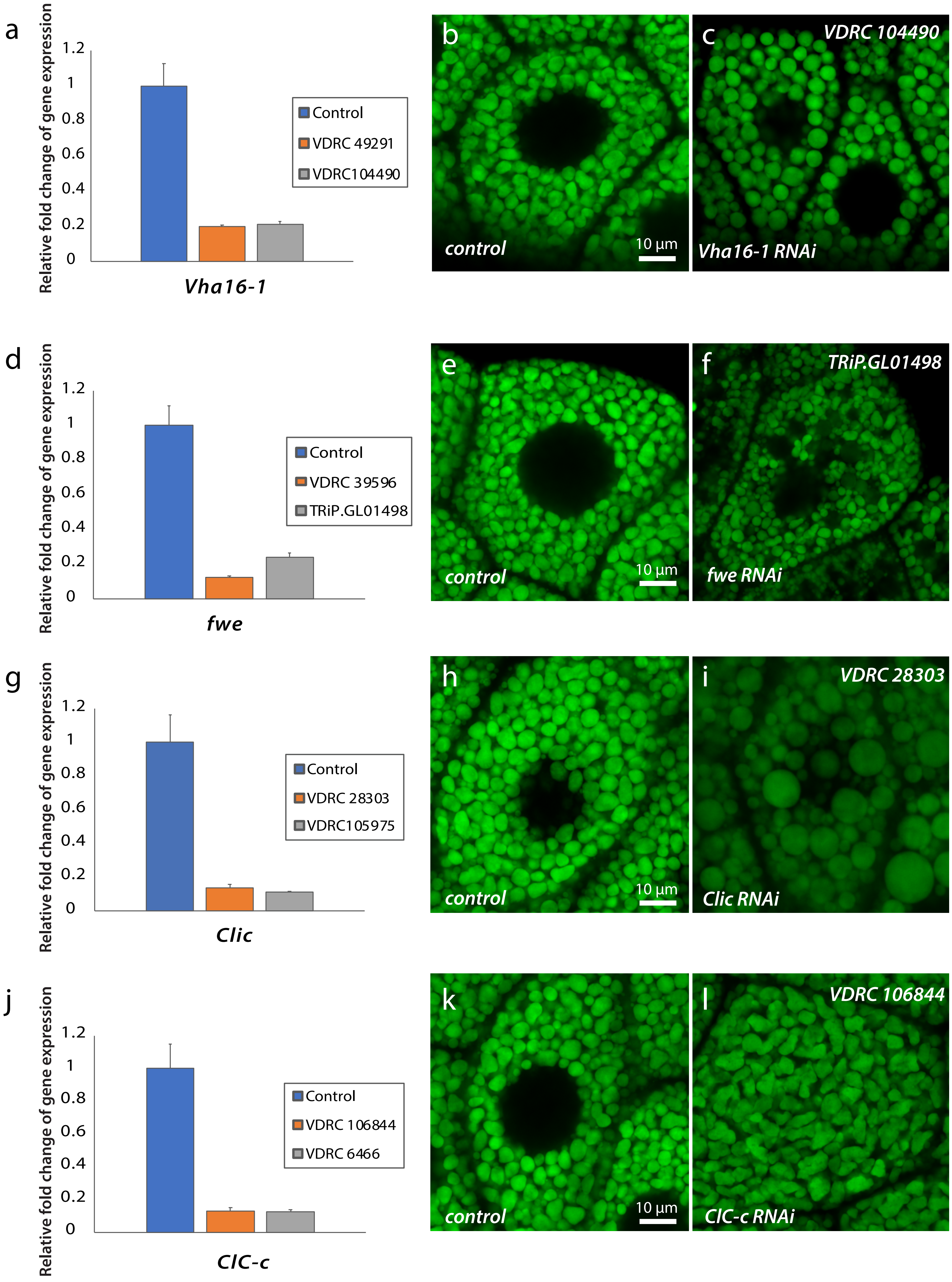

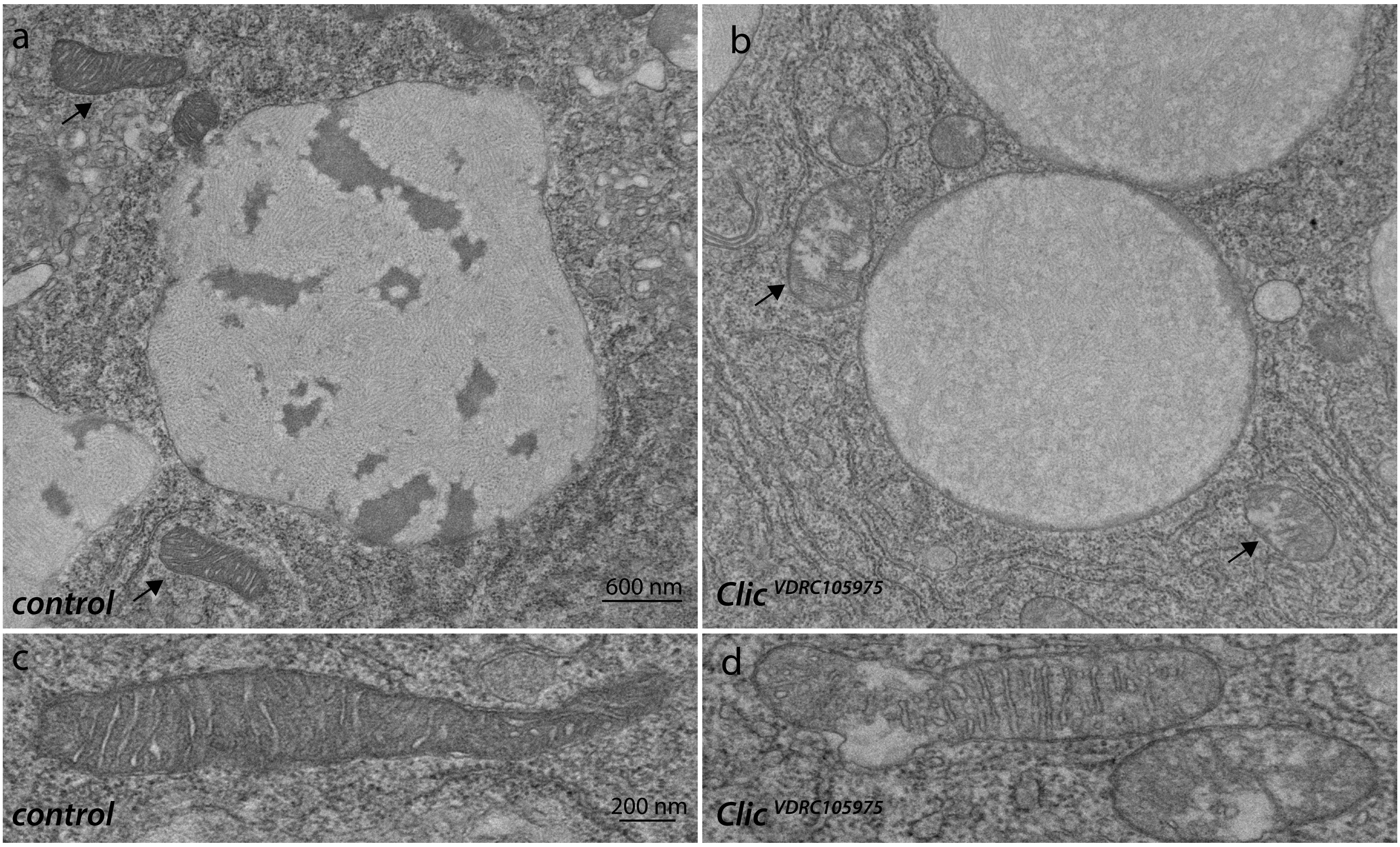
